# Latitudinal patterns of flowering phenology of three widespread tropical species from public media and data repositories

**DOI:** 10.1101/2023.08.18.553809

**Authors:** Bhavya Kriti, Sidharth Sankaran, Geetha Ramaswami

**Affiliations:** St. Xavier’s College, Ranchi, Jharkhand, India; Poorna Learning Centre, Bengaluru, India; Nature Conservation Foundation, Mysuru, India

**Keywords:** citizen science, latitudinal patterns, media repositories, tree phenology, tree seasonality, SeasonWatch

## Abstract

Plant phenology encompasses the study of periodic events in biological life cycles of plants and the influence of seasonal changes in weather and other environmental factors. In order to understand the role of the environment and the changing climate on the life cycle of plants, long-term records are required to establish baselines against which significant future deviations can be discerned. This study aims to analyze the spatial flowering pattern of the plant species -*Bombax ceiba, Butea monosperma* and *Cassia fistula* across India, through images uploaded on public media repositories like citizen science portals and social media. Citizen science is rapidly emerging as a reliable source of supplementary scientific information, and can be utilized to understand broad patterns of tree behavior. We found that for all three species, flowering period is longer, and begins earlier in southern latitudes as compared to the northern latitudes. We conclude that publicly available images of flowering trees and citizen science data can be used to supplement existing information on tree phenology across India.

## 1. Introduction

Phenology is the study of the timing of the specific life history events of a species, particularly in relation to cyclic seasonal changes (Raju *et al*. 2005; Jain *et al*. 2009; Sindhi and Bairwa 2010). This timing is an important factor in plant reproductive cycles and hence important to the population dynamics and persistence of a species (Jain *et al*. 2009; Sindhi and Bairwa 2010). Phenological events in mature plants include leaf-emergence, flowering, vegetative growth, fruiting, and leaf fall (Rathcke *et al*. 1985). Environmental variables such as temperature,precipitation, photoperiod, soil moisture, humidity, and soil nutrient content have been shown to influence plant phenology (Abu-Asab *et al*. 2001; Kudo and Ida 2013; Renner and Zohner 2018), and the same species may also show variable phenology depending on spatial factors such as latitude and altitude (Chaudhary and Khadabadi 2012). Of these factors, temperature and precipitation are likely to become highly variable due to global-warming induced climate change, in turn affecting plant phenology, typically observed as changes in the onset of a phenophase from a historical baseline (Walther *et al*. 2002; Cleland 2007; Gordo and Sanz 2010; Numata *et al*. 2022). Variability in environmental factors is likely to be exacerbated by anthropogenic changes such as urbanization, contributing to localized ‘urban heat islands’ or altered soil moisture due to artificial irrigation (Neil and Wu 2006). Climate change impacts on plant phenology are better-studied in temperate regions - where plants respond to lengthening growing seasons through advancing vegetative and reproductive phenology such as budburst and flowering (Badeck *et al*. 2004; Gordo and Sanz 2010). In recent years, due to climate change, seasonal climatic variations have begun to differ from their normal patterns (Rathcke *et al*. 1985). This could cause variations in the flowering dates (Renner and Zohner 2018; Numata *et al*. 2022) and reproductive cycles of many tree species which could in turn affect a whole host of animal species that depend on these trees to provide food and habitats. Plant phenology is linked to the life history patterns of consumers that feed on these species as well as their interactions with each other (Rathcke *et al*. 1985; Raju *et al*. 2005; Burli at al. 2007; Jain *et al*. 2009; Sindhi and Bairwa 2010). Consequently, changes in plant-pollinator relationships – where change in the timing of cyclic life events could lead to phenological mismatch-can negatively impact the interacting species (Renner and Zohner 2018). For example, early onset of flowering in the pan-Asian tuberous plant species *Corydalis ambigua* is associated with a lowered rate of pollination by bumblebees and therefore in lower seed production (Kudo and Ida 2013). Thus knowledge of the phenology of a species is important as it can be used to predict how the species may respond to climate change and the larger ecological implications at an ecosystem level.

India has a very wide variation in latitudes, extending between 8°4’N and 37°6’S and the climate varies accordingly – with very little temperature seasonality in the south and high temperature seasonality in the north, west and eastern regions of the country (Attri and Tyagi 2010). A large part of the Indian subcontinent is thus dominated by widespread tropical species. We hypothesise that species with ranges spread across latitudes, will show variable phenological behaviour as one can expect that the optimal temperature, and other environmental conditions such as day-length and precipitation, affecting phenology in these species would also vary considerably with latitude. The change in the timing of vegetative and reproductive phenology in tropical tree species is poorly understood due to lack of historical documentation.

Information on species-level phenology is often difficult to find outside of regional floras and other technical sources. Ideally, population-level replicates of species should be considered for such analyses. However, in the absence of intensively sampled data, one can derive information from large online public repositories of species images (Barve 2014) or from citizen science projects such as SeasonWatch (Ramaswami *et al*. 2021). SeasonWatch is a pan-india project monitoring the phenology of common tropical tree species wherein, citizen scientists make quantitative assessment of tree phenophases on a weekly basis, observing registered trees again and again with the aim of understanding long-term changes in tree seasonality (Ramaswami *et al*. 2021).

In this study, we explore the effect of latitude on the flowering phenology of three wide-ranging tree species *Bombax ceiba L*., *Butea monosperma (Lam*.*) Taub*, and *Cassia fistula L. Bombax ceiba* (Red Silk Cotton tree), belonging to the family Malvaceae, is a large deciduous tree, easily identifiable by its spiny bark and large pentamerous red flowers. The tree typically blooms before the onset of summer and attracts a large diversity of pollinators such as birds, bats and bees (Raju A. J. S., 2005). *Butea monosperma* (Flame-of-the-Forest tree), belonging to the family Fabaceae, is found in dry-deciduous, seasonal forest habitats of India. The bright red flowers of *Butea monosperma* are frequented by as many as seven species of birds, belonging to six families and one species in particular, the purple sunbird (*Nectarinia asiatica*), is an effective pollinator (Tandon *et al*. 2003). *Cassia fistula* (Indian Laburnum tree), also a Fabaceae member, has a wide geographical range, is native to the Indian subcontinent, and produces bright yellow flowers in large, pendulous inflorescences. *Xylocopa* bees (Carpenter bees) are known to be the most effective pollinator of *Cassia fistula* as they show high specificity towards this particular plant species (Kallur, 1993). All three species find use in local cuisine, medicine, and cultural practices, and because of their bright, showy flowers are likely to be identified accurately, observed frequently, and documented widely in media repositories and citizen science.

## 2. Methods

We used two data sources for understanding phenological patterns of these three species: 1) images of flowering trees available publicly on social media and other image repositories, and 2) phenology information submitted by citizen scientists to SeasonWatch. We expected latitudinal patterns in flowering to be discernible in both datasets, with the date of observation showing correlation with the latitude at which the phenophase was observed. We expected patterns observed through one-time phenological observations (images) to be supplemented long-term repeated observations from SeasonWatch to infer advantages of either method in assessing tree phenology.

In the first data source (dataset 1 hereon) - we collated images uploaded for *Bombax ceiba, Butea monosperma* and *Cassia fistula* on social media platforms (Facebook, Twitter, Instagram); image-based citizen science portals (Indian Biodiversity Portal, iNaturalist), and digital photo repositories (Flickr) between 1 January 2018 and 30 November 2021. All these platforms allow contributors to provide metadata on species ID, location and date of observation of a photograph. Images were searched through keywords such as scientific and common names of the species in English and other Indian languages, and through hashtags where appropriate. Images which included accurately identified plants with recognisable phenophases, and had reliable metadata such as the date of observation/image, and location were considered appropriate for the assessment of the relationship between tree phenology and latitude. We noted the presence of 4 phenophases - young leaves, mature leaves, unripe fruits and ripe fruits, in addition to the presence of 2 stages of flowers - buds and open flowers. Only images with the two flowering phenophases were considered for further assessment. We found 276 images of *Bombax ceiba*, 350 images of *Butea monosperma*, and 247 images of *Cassia fistula* in which the flowering phenology was clearly identifiable. We assessed flowering patterns visually by plotting the day of the year on which an image was taken, against the latitude of observation.

The second data source was the SeasonWatch database^19^ which was queried for the three species of interest. Citizen scientists register trees with the project and record phenophases (same as in dataset 1) as ‘none’, ‘few’, or ‘many’ based on the proportion of the tree canopy covered by the phenophase. We assessed flowering visually by plotting the day of the year on which an observation of ‘many’ flowers was made (indicating peak flowering), against the latitude of the observation. While both datasets allow for looking at latitudinal patterns in flowering, SeasonWatch data additionally allows for observing flowering patterns across the same individuals over time. We used repeated observations from 762 *Bombax ceiba*, 371 *Butea monosperma*, and 3329 *Cassia fistula* trees between 1 January 2018 and 30 November 2021 in the SeasonWatch database to assess the date of observation of flowering against the latitude of observation.

## 3. Result and Discussion

We did not combine information from dataset 1 and SeasonWatch as the raw information was collected in a non-comparable way (i.e. image and quantity of phenophase). In dataset 1, images of flowering *Bombax ceiba* trees appear between days 330 of the previous year and day 90 of the subsequent year, indicating December - March as the flowering season for this species (Figure 1a). In more southern latitudes (for example: 8.62°N-16.7°N) flowering trees were photographed from December - February while in more northern latitudes flowering trees were photographed in February-March, sometimes extending till May (Figure 1a). More continuous data from SeasonWatch indicates that at lower latitudes a few trees may be flowering throughout the year, while at latitudes above 9 degrees, most trees flowered between the 300th day of one year and the 150th day of the next (Figure 2a). A slightly more pronounced effect of latitude is seen for flowering in dataset 1 *Butea monosperma*, with most images appearing across latitudes between December (day 350) and March (day 90), sometimes extending till April or May (Figure 1b). SeasonWatch data indicate that trees have shown ‘many’ flowers as early as mid-year in latitudes below 13 degrees, but in more northern latitudes peak flowering occurred within the first 100 days of the year (Figure 2b). Images of flowering *Cassia fistula* trees occurred in dataset 1 almost throughout the year. Most images of flowering *C. fistula* trees across the 3.5 years were taken between the 50th and 200th day of the year (i.e. between end-February and July, Figure 1c). In SeasonWatch dataset, *C. fistula* trees were observed to flower almost throughout the year between 8- and 13-degrees N latitude, while flowering peaked more seasonally at higher latitudes (Figure 2c).

**Figure 1.**
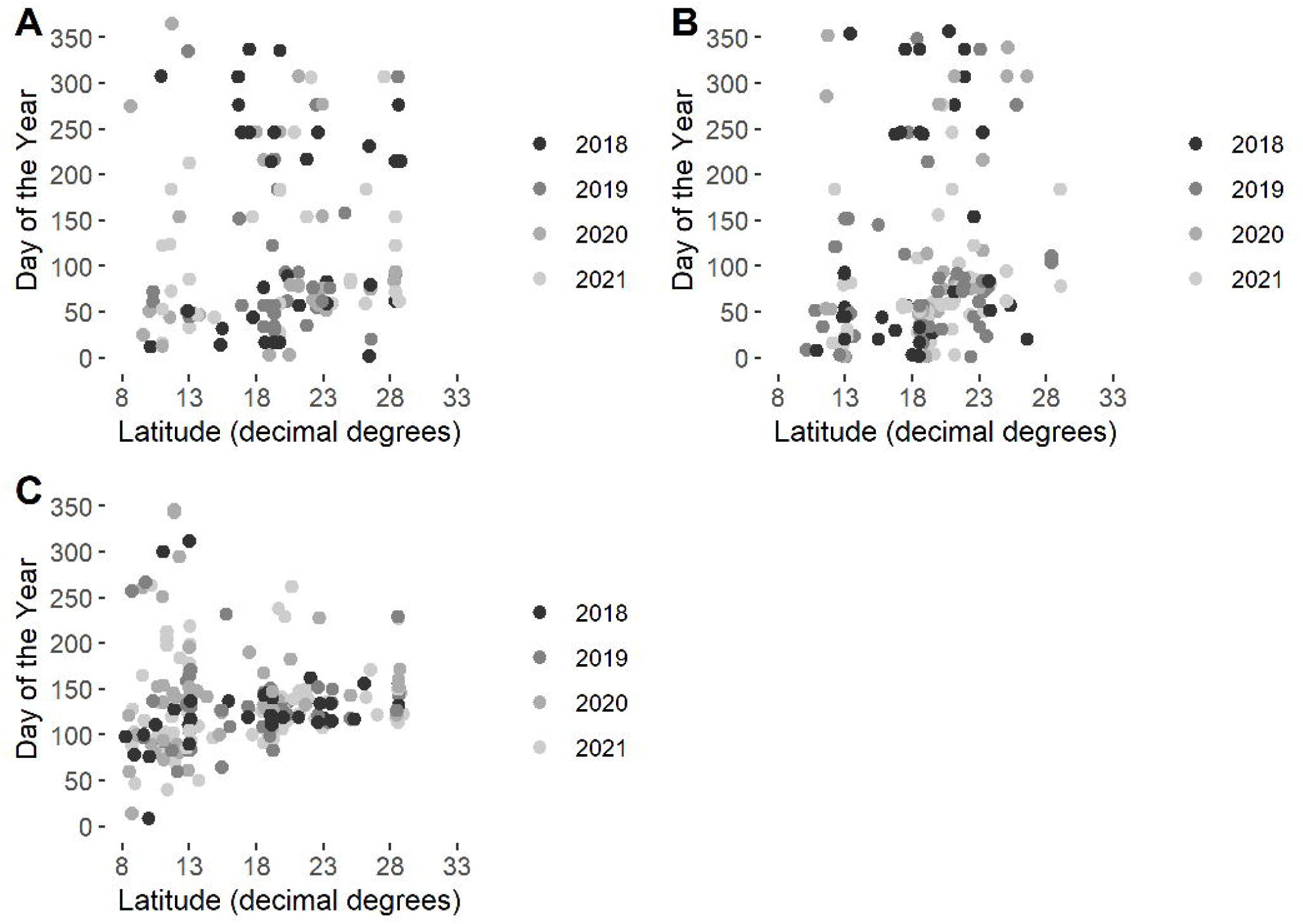
Phenology patterns from dataset 1 - date of upload of images of a flowering tree of A) *Bombax ceiba*, B) *Butea monosperma*, and C) *Cassia fistula*, against latitude at which the image was taken.

**Figure 2.**
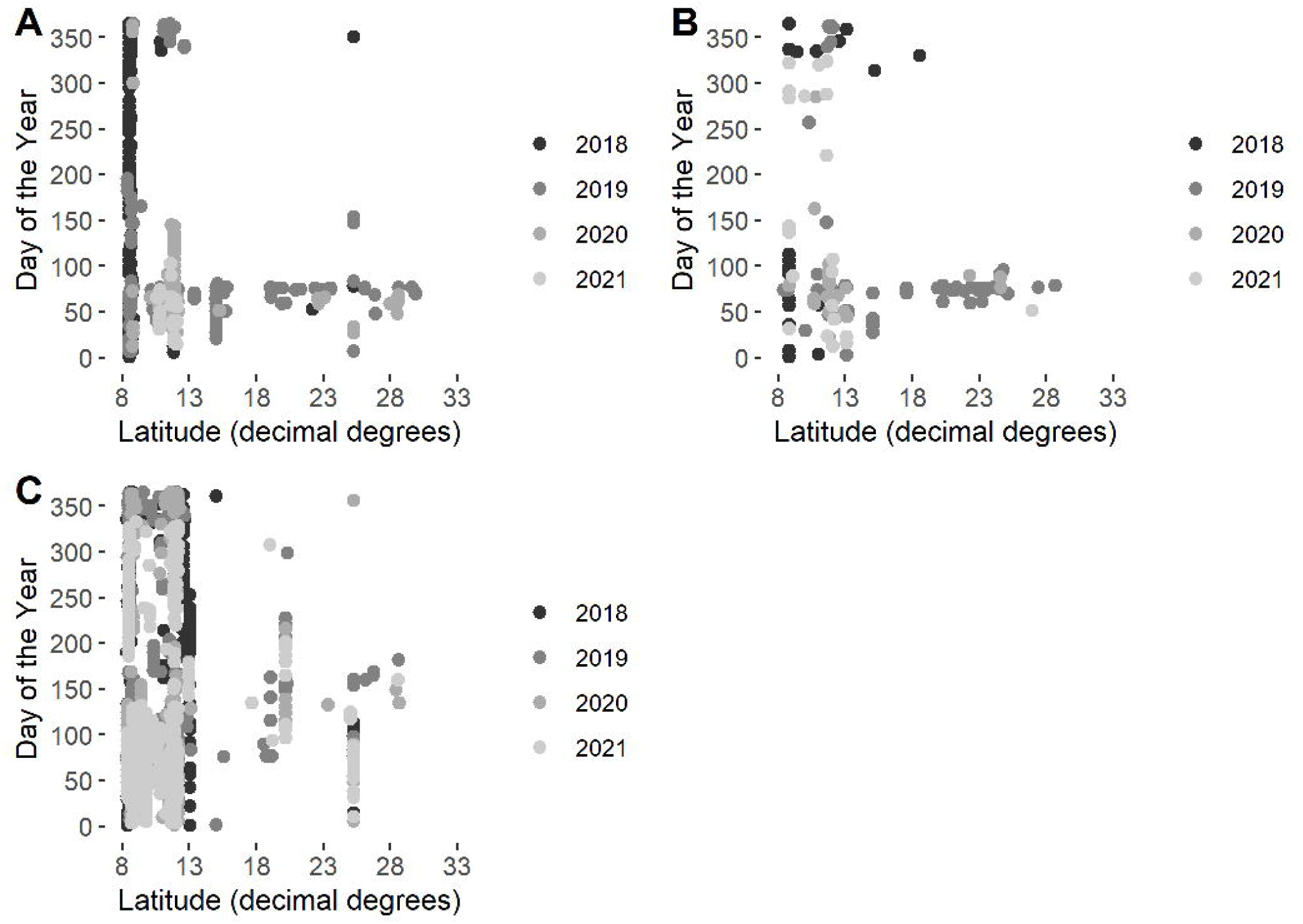
Phenology patterns from SeasonWatch - date of observation of a tree in peak flowering of A) *Bombax ceiba*, B) *Butea monosperma*, and C) *Cassia fistula*, against latitude at which the observation was made.

Across both datasets, the probabiltity of finding flowering phenology dates were consistently more in the first quarter of the year for *Butea monosperma* and *Bombax ceiba*, but staggered by latitude for *C. fistula*, when data was pooled across all years (Supplementary figure 1 and Supplementary figure 2). In the SeasonWatch dataset, C. Fistula appears to be more variable than the other two species, whereas in public media repositories, all three species show similar amount of variability in dates of flowering. On comparing the patterns from the two datasets, we infer that continued monitoring of individual trees allows for a better understanding of the flowering season of trees across latitudes - as individual-level variation is better captured through repeated observations. We do not have adequate understanding of what environmental factors could be driving phenological variation in these species across latitudes from this study alone, and speculate that as seen in other species, temperature and precipitation may be key drivers. If these species have a temperature threshold that triggers flowering then one could expect the flowering season to increase in duration across latitudes over the next few years as global temperatures increase (Rathcke *et al*. 1985). Citizen science data on flowering and leaf-out patterns elsewhere have demonstrated early onset of vegetative and reproductive phenophases with increasing mean spring temperatures (Fucillo Battle *et al* 2022). However, the wide variation in existing phenological strategies of tropical species such as the ones examined in this study, are likely to make temperature-phenology relationships less discernible than temperate species.

Images in dataset 1 are, at best, one-time snapshot samples of single individuals and may not represent the entire variation in the phenological behaviour of the study species. Furthermore, contributor behaviour is likely to affect what images are posted publicly, and which May in-turn bias the data available on these platforms. For instance, contributors may be more likely to post photos of clearly visible flowers and of trees in full bloom, than any given instance of flowering, particularly in the species examined. This may create a bias towards a more visually identifiable stage of the flowering phenopahse, than the entire duration of the phenology. Contributor behaviour may also be affected by other factors such as access to smart phones or cameras to take pictures, or access to public groups and repositories, because of technological or linguistic constraints. Therefore, dataset 1, may have lower utility in understanding future changes in tree phenology with changing climate as compared to SeasonWatch data that has multiple observations of individuals in the same locations over time. However, in order to utilize citizen science data effectively, high data quality needs to be ensured. Phenology citizen science assessed elsewhere has found disparities between citizen contributed data and actual observations (MacEnzie *et al*. 2017). Although a similar assessment, we conclude that both approaches to observing tree phenology can be used to supplement existing phenological information on the species, or to arrive at the baseline patterns of phenology to aid in designing sampling strategies for future studies.

## Acknowledgements

Bhavya Kriti would like to thank the Indian Academy of Science for supporting this study. SeasonWatch is supported by the Wipro Foundation. We would like to thank the hundreds of citizen scientists who contributed the data to the SeasonWatch database and facilitated this comparative study.

## Conflict of Interest

The authors do not have any conflict of interest to report.

## Notes

### Competing Interest Statement

The authors have declared no competing interest.

### Summary of Updates

Supplementary images have been added for clarity of the results as well as certain changes in the literature has been made upon suggestion by reviewers.

