## Supplementary figures and images for "Latitudinal patterns of flowering phenology of three widespread tropical species from public media and data repositories"

### Supplementary figure 1

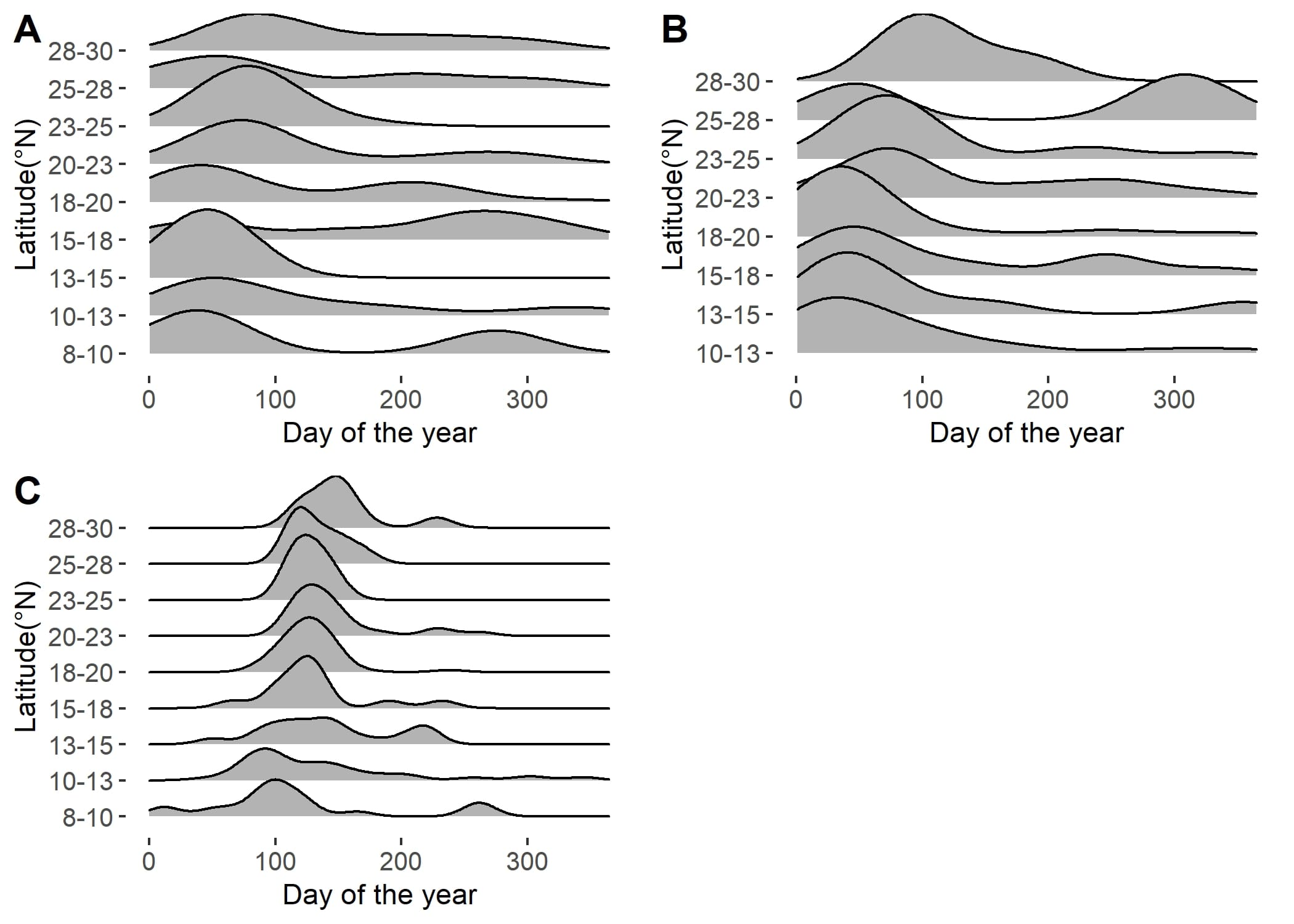

### Supplementary figure 2

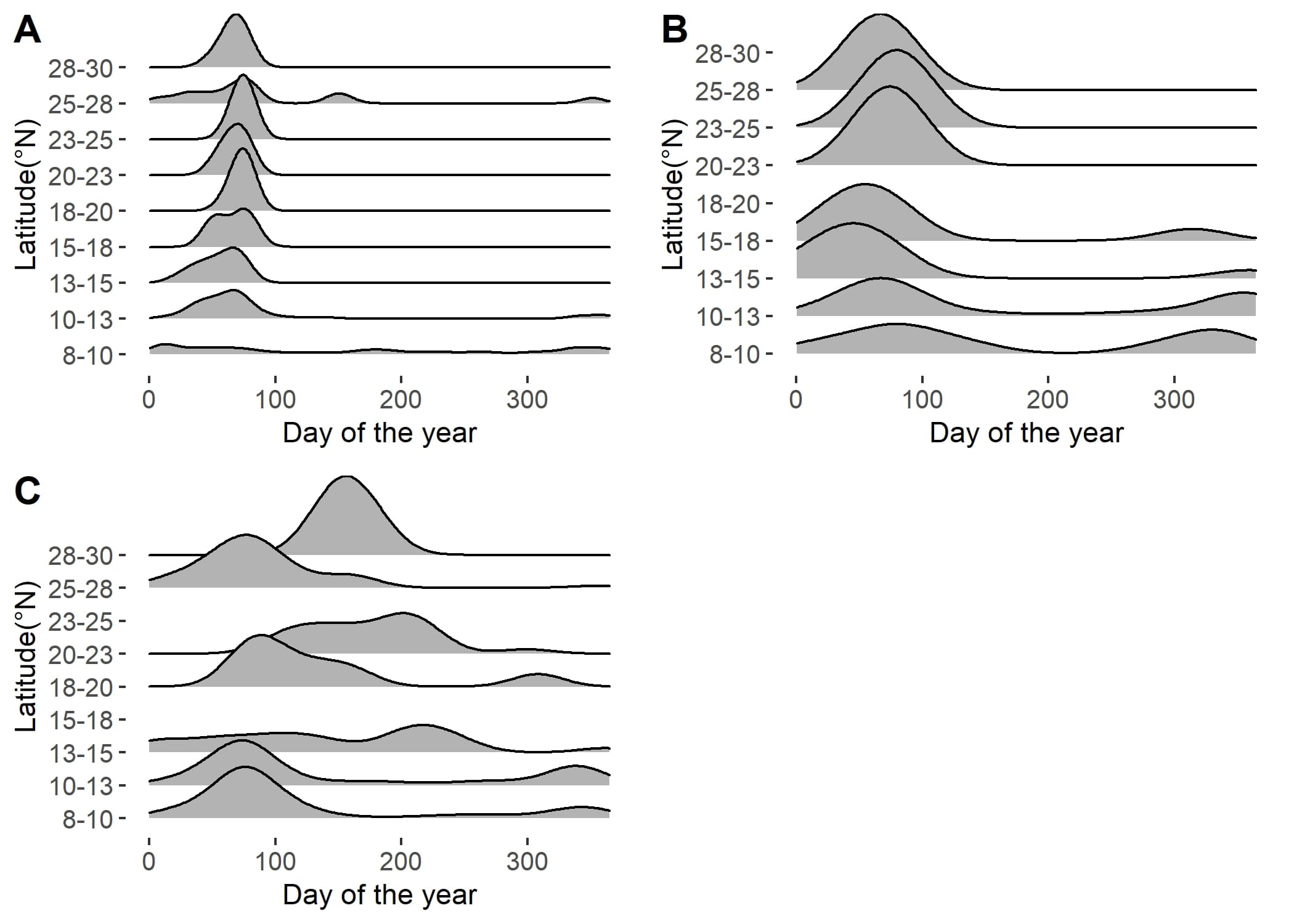
